# Deep Learning from Phylogenies for Diversification Analyses

**DOI:** 10.1101/2022.09.27.509667

**Authors:** Sophia Lambert, Jakub Voznica, Hélène Morlon

## Abstract

Birth-death models are widely used in combination with species phylogenies to study past diversification dynamics. Current inference approaches typically rely on likelihood-based methods. These methods are not generalizable, as a new likelihood formula must be established each time a new model is proposed; for some models such formula is not even tractable. Deep learning can bring solutions in such situations, as deep neural networks can be trained to learn the relation between simulations and parameter values as a regression problem. In this paper, we adapt a recently developed deep learning method from pathogen phylodynamics to the case of diversification inference, and we extend its applicability to the case of the inference of state-dependent diversification models from phylogenies associated with trait data. We demonstrate the accuracy and time efficiency of the approach for the time constant homogeneous birth-death model and the Binary-State Speciation and Extinction model. Finally, we illustrate the use of the proposed inference machinery by reanalyzing a phylogeny of primates and their associated ecological role as seed dispersers. Deep learning inference provides at least the same accuracy as likelihood-based inference while being faster by several orders of magnitude, offering a promising new inference approach for deployment of future models in the field.

## INTRODUCTION

Phylogenetic approaches for studying species origination and extinction dynamics over deep time rely on the statistical adjustment of stochastic birth-death models (Kendall 1948) to dated phylogenetic trees representing the evolutionary relatedness of species and the dating of their divergence times (Stadler 2013; Morlon 2014; Harmon 2019). An increasing amount of such phylogenetic data have become available, and has been accompanied by complexification of diversification models. These models include homogeneous rate models where speciation and extinction rates are identical across lineages at any given time, and range from simple time-constant (Nee et al. 1994) to time-dependent (Morlon et al. 2011; Stadler 2011; May et al. 2016), environment-dependent (Condamine et al. 2013), and diversity-dependent (Etienne et al. 2012) models. Diversification models also include heterogeneous rate models (Alfaro et al. 2009; Morlon et al. 2011; Rabosky 2014; Höhna et al. 2019; Maliet et al. 2019; Barido-Sottani et al. 2020; Laudanno et al. 2020), with the class of State-dependent Speciation and Extinction (SSE) models that links rate heterogeneity to specific characteristics of the species (Maddison et al. 2007; FitzJohn 2010; Goldberg et al. 2011; Fitzjohn 2012; Beaulieu and O’Meara 2016; Herrera-Alsina et al. 2019; Vasconcelos et al. 2022).

The parameters of interest of these models – mainly speciation and extinction rates – are traditionally inferred using likelihood-based techniques – Maximum likelihood or Bayesian inference. Maximum likelihood consists in finding the parameters that maximize the probability of observing the data (here the phylogeny, or the phylogeny and associated trait data in the case of SSE models); Bayesian inference uses this probability along with *a priori* information on the parameters of interest to explore the *posterior* probability distribution of the parameters knowing the data. Numerous studies have used these inference approaches to estimate speciation and extinction rates over geological times and across the Tree of Life, to investigate the processes modulating these diversification dynamics (Stadler 2011; Etienne et al. 2012; Pyron and Wiens 2013; Höhna 2014; Rolland et al. 2014; Gubry-Rangin et al. 2015; Rabosky et al. 2018; Condamine et al. 2019; Stone and Wolfe 2021). While powerful, likelihood-based inference techniques are limited by potential intractability issues for complex diversification models, and by computational cost on increasingly large phylogenetic data (Hinchliff et al. 2015). Indeed, complex models do not always have a closed-form solution and/or the likelihood cannot always be evaluated in a reasonable amount of time. This renders the application of likelihood-based methods to complex models and species-rich groups (*e*.*g*., insects, micro-eukaryotes and prokaryotes) difficult. As a result, there are several models of diversification in the literature which behavior has been studied with simulations but that lack a proper inference machinery (McPeek 2008; Aristide and Morlon 2019; Hagen et al. 2021). Methods based on Expectation Maximization (EM) algorithms (Dempster et al. 1977; Richter et al. 2020), data augmentation (Maliet and Morlon 2022), or composite likelihoods (Lindsay 1988; Varin et al. 2021)) can overcome some of these limitations (Raynal 2019), yet they still rely on likelihood formulae.

An alternative is the use of likelihood-free inference techniques, such as Approximate Bayesian Computation (ABC (Beaumont et al. 2002; Marin et al. 2012; Sisson et al. 2018)). In its most basic form, ABC relies on generating artificial data by simulating the process of interest along a given parameter range and compressing the data by computing summary statistics on these simulations to enable the comparison between simulated and observed summary statistics. This comparison is done by computing a distance and evaluating if this distance is sufficiently small to accept the simulated data using a chosen tolerance threshold (rejection-based approach). ABC has been useful to fit complex models in various fields, including phylogenetic diversification analyses (Bokma 2010; Janzen et al. 2015; Janzen and Etienne 2016). However, there are several limitations of ABC, including its reliance on the choice of summary statistics that should resume the information contained in data in a low number of metrics, the choice of the distance metric to compare the observed and simulated data and the choice of the tolerance threshold. While it is possible to evaluate and minimize the influence of these choices on parameter inference (Sisson et al. 2007; Beaumont et al. 2009; Blum and François 2010; Del Moral et al. 2012; Blum et al. 2013; Prangle 2017), an adjustment needs to be performed each time a new model is developed. In particular, new summary statistics must be designed to convey information relative to the problem at hand.

Deep learning offers an alternative likelihood-free inference technique. Deep learning (Goodfellow et al. 2016) is a subfield of machine learning where highly flexible statistical learning functions based on neural networks (NNs) are used to learn regression (such as parameter estimation) or classification (such as model selection) problems. The term ‘deep’ is conventionally associated with a neural network that takes raw data as input values and extracts patterns from this low level representation thus creating its own ‘summary statistics’ or high-level features, without the need of designing those. This definition is the one used in the present article. Applying a regression task with deep learning (learning model parameter values from simulations) is increasingly used in several fields including population genetics (Sheehan and Song 2016; Sanchez et al. 2020; Avecilla et al. 2022), phylogenetic reconstruction (Nesterenko et al. 2022), macroecology (Andermann et al. 2022) and physiology (Kroll et al. 2021). In macroevolution, an early progress of using machine learning (Bokma 2006) consisted in training an artificial neural network on the axes of a principal component analysis of phylogenetic branching times to infer speciation and extinction rates. A similar framework was then used for the inference of rates of phenotypic evolution (Bokma 2010). While promising, this approach was not developed further by the community.

A step forward was recently taken by Voznica et al. (2022), who developed a deep learning approach for the statistical inference of birth-death models from phylogenies in the context of pathogen phylodynamics. The authors developed a tree representation, the Complete Bijective Ladderized (CBLV) tree representation, which applies to non-ultrametric trees representing the evolutionary relationship between pathogen sequences sampled at different dates. The CBLV representation proved to be efficient for model selection and inference of transmission dynamics when combined with Convolutional Neural Networks (CNN) (LeCun et al. 1998), where it yielded accuracy at least comparable to gold-standard Bayesian approaches. Voznica et al. (2022) also combined an extensive set of Summary Statistics with Feed-Forward Neural Networks (FFNN-SS) that yielded similar results. To our knowledge, comparable attempts to use deep learning to infer diversification dynamics from species phylogenies do not exist.

Here, we adapt the approach of Voznica et al. (2022) to birth-death diversification models used to infer diversification dynamics from phylogenies of extant species (and also potentially associated trait data). The phylogenetic data are different from the one in Voznica et al. (2022). First, the evolutionary trees are different as, if no data from fossils is included, the species are all sampled at present time and the reconstructed phylogenies are thus ultrametric, that is the tips are all at the same distance to the root. Second, we allow for possibility to include trait data associated to the tips. We begin by adapting the CBLV tree representation from Voznica et al. (2022) to ultrametric phylogenies and to ultrametric phylogenies with tip state data, in the form of ‘Complete Diversity-reordered Vector’ (CDV). We then assess the performance of deep learning inference in comparison to maximum likelihood estimation (MLE) by using the simple homogeneous time-constant birth-death (BD) model (Nee et al. 1994), for which a closed-form expression of the likelihood exists, and the binary-state speciation and extinction (BiSSE) model, for which the likelihood is approximated by solving Ordinary Differential Equations (Maddison et al. 2007). Finally, we illustrate the approach by applying our trained neural network for BiSSE to an empirical phylogeny of 273 primates (Fabre et al. 2009) and their associated interaction type (mutualistic or antagonistic) with plants (Gómez and Verdú 2012).

## METHODS

Our main goal is to develop a compact and exhaustive representation of the raw data into a matrix (the CDV), and to test the performance of a CNN combined with this representation for parameter inference (thereafter referred to as the CNN-CDV approach). We train the neural networks with data simulated under the birth-death diversification processes. For the time-homogeneous birth-death model (BD), we compare the performance of CNN-CDV to FFNN combined with a series of summary statistics (FFNN-SS), FFNN combined with the CDV representation (FFNN-CDV), and the maximum likelihood estimation (MLE) approach. For the BiSSE model, we compare the performance of CNN-CDV to FFNN-CDV and MLE. Codes used to perform the simulations, encode the phylogenetic data into the CDV representation, train the neural networks, and use the trained networks on simulated or empirical data for parameter inference, are available on GitHub (https://github.com/JakubVoz/deeptimelearning).

### TREE REPRESENTATIONS

#### Full Tree Representation: Compact Diversity-reordered Vector (CDV)

We adapted the Compact Bijective Ladderized Vector (CBLV) representation used in Voznica et al. (2022), originally intended for non-ultrametric trees, to ultrametric trees and to ultrametric trees with information on tip states. The encoding proceeds in the following steps (**Fig. 1)**: 1] internal node reordering, 2] inorder tree traversal (Cormen 2009) and creation of vector representation, 3] completion to the maximum tree-size in simulations, 4] addition of tree height and sampling probability to the vector representation.

**Figure 1:**
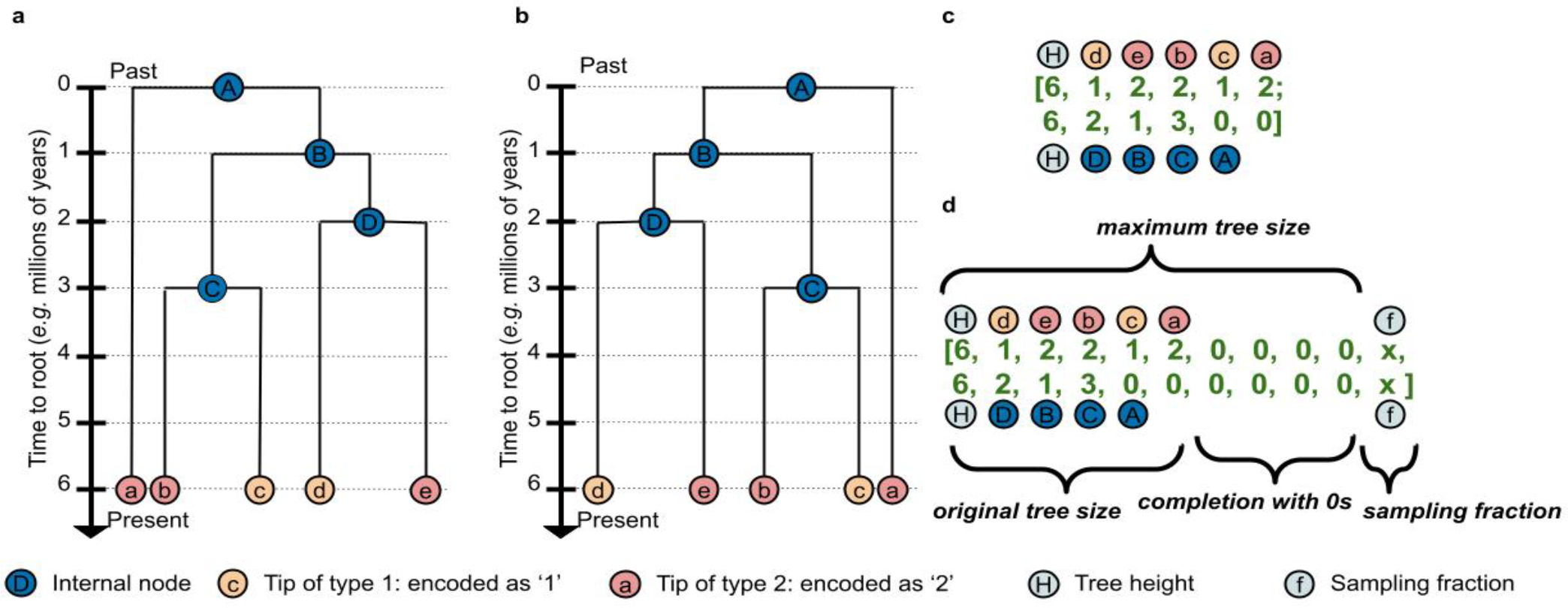
Representation of ultrametric trees with tip state data. Illustration of the encoding algorithm for ‘Compact Diversity-ordered Vector’ (CDV) on a tree with 5 tips. ***(a)*** Ultrametric tree with tip state information (here there are two possible states for each tip, either state 1 or state 2, represented in orange and pink respectively). ***(b)*** The tree is reordered following a diversity criterion: for each internal node, the sum of the branch lengths of the descending tree is computed and the internal node with higher sum is rotated on the left. ***(c)*** We then create a 2-rows matrix filled by visiting the tree according to a tree inorder traversal algorithm, with tips represented on the top row and internal nodes on the bottom row. Tips are assigned their encoded trait state (‘1’ for state 1 and ‘2’ for state 2) and internal nodes their distance to the root. We add a first column with tree height. ***(d)*** Finally, we complete the matrix with zeroes, so that its size is the one of the largest simulated tree and we add a column with the value of the sampling fraction. To obtain the representation of an ultrametric tree without tip state data, we do not include the first row on tip state information.

Before encoding, we rescaled the trees to unit average branch length and the rate parameters (*e*.*g*. diversification rate(s), turnover rate(s) *etc*.) accordingly. The trained neural networks should thus apply to trees with any time scale. The criterion used in Voznica et al. (2022) for tree reordering was based on the ladderization, where each internal node is rotated so that the branch supporting the most recent tip is on the left. As this cannot apply to ultrametric trees where all tips are sampled at the same time, we used a diversity criterion: for each internal node we compute the sum of the branch lengths of the descending tree (i.e. phylogenetic diversity) and the branch with highest sum is shifted to the left. Next, we perform a tree inorder traversal: the reordered phylogeny is traversed by recursively starting with the left subtree and for each visited internal node, its distance to the root is added to a vector (Cormen 2009). For trees with tip data, we create a second vector with information on the tip data, while visiting tips during the same traversal. For binary tip data (as obtained by simulating BiSSE for example), we use the values 1 and 2 to distinguish the two states. These two vectors are then combined into a matrix. We then add a first column with the value of the tree height. We name this representation a Compact Diversity-ordered Vector (CDV). This representation could be easily extended to account for information on non-binary traits, for example using one hot encoding, which consists in encoding a qualitative variable of x states into x rows with 1 representing the presence of state and 0 its absence. Similarly, multiple traits, including quantitative ones, can be encoded by stacking additional rows to the matrix. Furthermore, we can imagine treating missing trait data by assigning a value of −1 in the CDV representation when the information is lacking.

The CDV representation is bijective (under mild assumptions, for example the absence of concomitant branching events), in the sense that we can unambiguously reconstruct any given tree from its representation, and compact: (x+1)\**n* entries for a tree with *n* tips (n-1 values for internal nodes and 1 on tree height) and x informational values on tips (x=0 in the absence of tip state information and x=1 for information on a single trait). The created vector (or matrix when tip state information is available) is completed with zeroes to obtain a representation of the same size as the largest phylogeny in the simulations. In order to account for potential missing extant species in the phylogeny, we add a last column with the value of the sampling fraction, computed as the ratio of the number of species represented in the phylogeny divided by the total number of extant species.

In order to assess whether the CDV representation with tip data allows to properly capture tip information, we compared results obtained with this representation to those obtained with a less informative representation (coined CDV-less) where we add only two values on tip type counts (*i*.*e*. the number of tips in each state) at the end of the row summarizing internal nodes.

#### Summary statistics representation

We used a set of 97 summary statistics (SS) representing trees (without associated trait data as we do not implement the FFNN-SS for the BiSSE model). The summary statistics were mainly based on those published in Saulnier et al. (2017) and in Voznica et al. (2022) (see the original papers for details) and they comprise:

- 8 summary statistics on tree topology
- 25 summary statistics on branch lengths
- 49 summary statistics on the Lineage-Through-Time (LTT) plot
- 14 summary statistics on consecutive internal branches
- 1 summary statistics on number of tips

We modified several statistics so that they apply to ultrametric trees. Instead of minimal and maximal tree height (the time of first and last sampled tips in non-ultrametric trees), we use the crown age. In Saulnier et al. (2017), there are several summary statistics defined on the LTT plot which consider the maximum number of the living lineages, the time of its occurrence and the slopes of the curves before and after this time. In ultrametric trees, the maximum number of the living lineages always appears at present (the moment of sampling tips) and thus such division of LTT plot is not possible. Instead, we divide the LTT plot into three equal parts and we measure the slope for each one, together with the ratios between the first and the second slope and between the second and the third slope. The computing time of these statistics grows linearly with tree size. We added the sampling fraction to these 97 measures, thus resulting in a vector of 98 scalars.

Finally, we reduced and centered the SS by subtracting the mean and scaling to unit variance, using the standard scaler from the scikit-learn package (Pedregosa et al. 2011) fitted to the training set.

### NEURAL NETWORKS: ARCHITECTURE AND TRAINING

A NN is organized in neural layers that in turn are organized in neurons (or ‘units’). In supervised learning, a NN can be trained to minimize the difference between an expected output (or target) and the predicted one, the measure of the difference being called a ‘loss function’. Here we used the mean absolute error as the loss function. A NN contains an input layer (by which the numerical values representing the data are passed to the following layer) and an output layer (in our case outputting parameter values), potentially separated by hidden layers. If there is at least one hidden layer, we talk about deep neural networks.

Feed-Forward Neural Networks (FFNNs) are one of the most basic forms of deep NNs. They are fully connected: for each neural layer, all neurons are connected to the neurons of the previous layer. The connections are characterized by trained bias and weights *i*.*e*. real values by which individual inputs of a given neuron are multiplied, as well as an activation function by which the summed input (input values multiplied by weights to which a bias value is added) is transformed. Here, we used the exponential linear function as activation function (Clevert et al. 2015). FFNNs typically work well on structured data, such as summary statistics, where the same summary statistics are at exactly the same entry in the input vector. On unstructured data (such as images or CDV) however, the number of parameters to train increases quadratically with the size of the input, and the information is scattered along the input vector, making FFNNs potentially less efficient.

Convolutional Neural Networks (CNNs) (LeCun et al. 1998) contain a convolutional part and a fully connected one. The convolutional part consists of convolutional and pooling layers and outputs a vector used as input of the fully connected part. The convolutional part aims at learning and extracting repeated patterns in the input, that are then combined for prediction in the fully connected part. Convolutional layers transform their input with several convolutional operations, each one specified by a kernel (or ‘filter’ or ‘feature detector’) whose parameters are trained. Each kernel transforms subparts or patches of the input by applying the convolutional operation and outputs a single value for each patch. We set the stride (by how much the kernel moves on the input when traversing it) to 1. A convolutional layer is specified by the number of kernels and their size (the size of their input), and outputs a ‘feature map’ (intermediate representation of transformed input). Pooling layers transform the resulting feature maps into smaller ones by taking the maximum or average of values subsampled in the map with a given window. CNNs typically work well with raw, low-level (vector and matrix) data such as images, videos or time-series recordings, by learning and extracting repeated patterns or ‘features’ through trained convolution functions and building from them their own high-level features (such as a set of summary statistics). They do not need prespecified feature input such as summary statistics. Each kernel learns and extracts one pattern in the data that can appear anywhere in the representation. In comparison to Voznica et al. (2022), increased kernel size in the first convolutional layer performed slightly better (see below; data not shown).

The training of a given network consists in iteratively changing the parameters of the NN (*e*.*g*. bias, weights) that minimize the loss function. This is performed by an optimization algorithm for stochastic gradient descent. Here, we used the Adam optimizer (Kingma and Ba 2015). The networks were trained on simulated data, the targets being the parameters of the diversification models (here BD and BiSSE).

Several training ‘tricks’ were developed for efficient and robust training in practice. We used a training set of 990.000 simulations split into subsets called batches, during the training. After measuring the loss on the whole batch, the trained parameters are updated with the optimizer to minimize the loss. Splitting the training set into batches of simulations enables to update the trained values more robustly (and moving into ‘right direction’ with respect to the minimal error). We set the batch size to 8,000 simulations. When the network parameters were updated on the whole training set (we talk about an ‘epoch’), the training starts again passing through the whole training set.

To prevent overfitting, we used a dropout of 0.5, which consists in shutting down randomly half of the neurons in the network during the training phase (Srivastava et al. 2014). We also used a technic called early stopping (Bengio 2012): at the end of each epoch, the loss is computed on a validation set (here, we used a validation set of 10.000 simulations), and the training is stopped when the loss on the validation set starts to increase. Typically, during the training of our networks, there will be hundreds of epochs before the training is stopped. The test set then enables to measure the true accuracy of the network. In our case it consisted of 500 simulations for the BD model and 10,000 simulations for BiSSE.

For the BD model, we used a CNN architecture on the CDV representation (referred to as CNN-CDV), a FFNN architecture on the CDV representation (FFNN-CDV), and a FFNN architecture on the summary statistics representation (FFNN-SS). For the BiSSE model, we used only the CNN-CDV and the FFNN-CDV, as applying the FFNN-SS would require deploying a new set of summary statistics accounting for tip data. We used the same FFNN architecture as in Voznica et al. (2022) in the case of both the FFNN-CDV and the FFNN-SS (see the original paper for details). The CNN architecture differed only by the size of the kernels (i.e. the size of their inputs) in the first layer; we used layers of size 5*2 and 5 for respectively the CDV and CDV-less as it performed slightly better than kernels of size 3*2 (respectively 3, data not shown). Also, unlike in Voznica et al. (2022), the function of the output layer was set to the exponential linear function (Clevert et al. 2015).

We implemented the NNs in Python 3.6 using the Tensorflow 1.5.0 (Abadi et al. 2016), Keras 2.2.4 (Chollet 2015) and scikit-learn 0.19.1 (Pedregosa et al. 2011) libraries.

### MACROEVOLUTIONARY MODELS AND SIMULATIONS

We assessed the performance of the NNs with two widely used models, the simple homogeneous, time constant birth-death (BD) model, and the binary-state speciation and extinction (BiSSE) model.

#### The Birth-Death (BD) Model

The BD model has a closed-form expression of its likelihood, which allows us to compare the performance of the Deep Learning and MLE approach in a “best case” scenario for MLE. Here, when referring to BD, we imply the homogeneous time-constant birth-death-sampling model (Yang and Rannala 1997; Stadler 2009), as we consider the possibility that some extant species are not represented in the phylogenies. In this model, new species originate with a constant speciation rate *λ* and go extinct with a constant extinction rate *µ*, typically expressed in number of events/lineage/Myr. At present each extant species is sampled with probability *f* (Bernoulli sampling scheme (Stadler 2009)). We assume *f* to be fixed, in which case *λ* and *µ* are identifiable.

We parametrized our simulations with the turnover (*ε = µ / λ*) and speciation rate (**Table 1**); the parameter values were sampled uniformly at random within parameter boundaries with standard Latin-hypercube sampling (McKay et al. 1979) using the Python PyDOE package. We performed the simulations with our own simulator, using a Gillespie algorithm (Gillespie 1977). Each simulation started with one lineage and ended when the number of living species reached 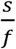, where *s* is the number of tips in the sampled phylogeny. We then sampled *s* species. We thus conditioned the simulations on the number of tips. We trained the NNs to learn *λ* and *ε*.

**Table 1:** Parameterization of the constant-rate birth-death model with incomplete sampling.

| Parameters | Symbols | Ranges |
| --- | --- | --- |
| **Turnover rate** | $\epsilon$ | **U(0.01,1)** |
| **Speciation rate** | $\lambda$ | **U(0.01,0.5)** |
| Tree size | s | U(200, 500) |
| Sampling fraction | f | U(0.01,1) |

#### The Binary State Speciation and Extinction (BiSSE) Model

The BiSSE model is the simplest state-dependent birth-death model: species are characterized by a binary state (1 or 2) which can influence their constant speciation (*λ*_*1*_ and *λ*_*2*_) and extinction rates (*µ*_*1*_ and *µ*_*2*_). Species can also transition anagenetically from one state to the other (with rates *q*_*12*_ and *q*_*21*_). As for the BD model, we can add to this diversification process a Bernoulli sampling scheme at present, which allows analyzing trees with missing extant species. We consider here a simple version of BiSSE with symmetrical transition rates *q=q*_*12*_*=q*_*21*_ as well as turnover rate and sampling probabilities at present (*ε* and *f*, respectively) shared across species irrespective of their state.

The BiSSE model allows us to illustrate the utility, and test the validity of the CDV representation with tip data. The likelihood of this model can be computed by solving Ordinary Differential Equations (ODEs), and recent efforts have been made to provide an efficient maximum-likelihood inference machinery for this model on large phylogenies (Louca and Pennell 2020).

We parameterized our simulations with the speciation rate associated to state 1 (λ_1_), the turnover rate (*ε*) and the ratios of λ_2_ and q_12_ relative to λ_1_. We sampled these parameters within a biologically realistic parameter space, given the literature on empirically inferred parameters (e.g. (Villarreal and Renner 2013; Williams et al. 2014; Gamisch 2016)) (**Table 2**). The parameter subspace was covered with standard Latin-hypercube sampling (McKay et al. 1979) using the Python PyDOE package. The simulations were performed using the R package castor 1.6.6 (Louca et al. 2018) and were conditioned on number of tips. We trained the NNs to learn λ_1_, λ_2_, q_12_ and *ε*.

**Table 2:**
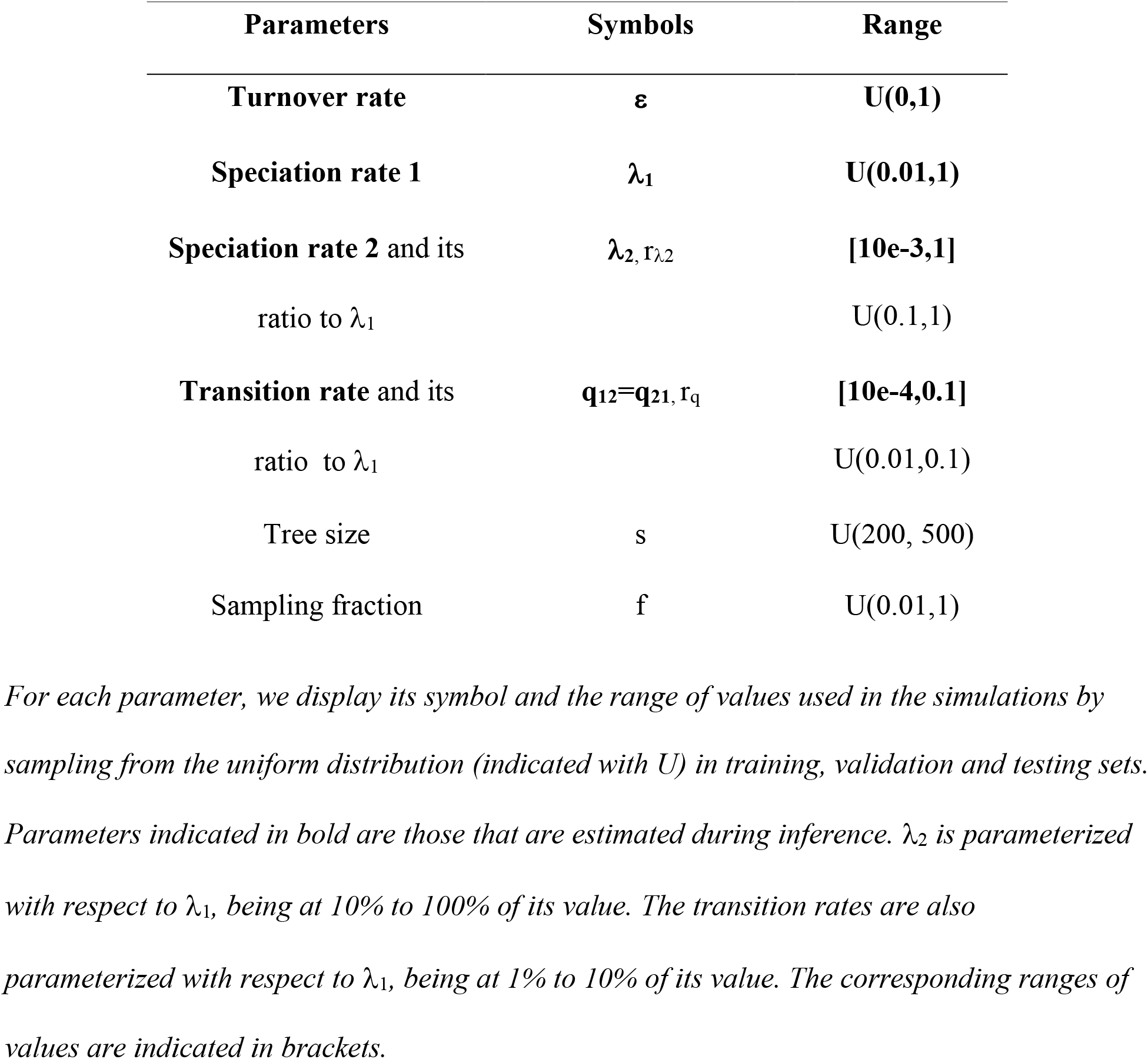
Parameterization of the Binary State Speciation and Extinction model with incomplete sampling.

| Parameters | Symbols | Range |
| --- | --- | --- |
| **Turnover rate** | $\epsilon$ | **U(0,1)** |
| **Speciation rate 1** | $\lambda_1$ | **U(0.01,1)** |
| **Speciation rate 2** and its<br>ratio to $\lambda_1$ | $\lambda_2, r_{\lambda_2}$ | **[10e-3,1]**<br>U(0.1,1) |
| **Transition rate** and its<br>ratio to $\lambda_1$ | $q_{12}=q_{21}, r_q$ | **[10e-4,0.1]**<br>U(0.01,0.1) |
| Tree size | s | U(200, 500) |
| Sampling fraction | f | U(0.01,1) |

### MAXIMUM LIKELIHOOD ESTIMATION

We compared the NN approaches with MLE. For the constant-rate BD, we used a MLE based on the Nelder–Mead optimization algorithm encoded within our custom function fitMLE_bdRho available at https://github.com/sophia-lambert/UDivEvo/tree/master/R. The function encodes a likelihood formula conditioned on the age of the phylogeny (here t_0_ = t_crown_) under a Bernoulli sampling scheme, and parametrized to infer the net diversification rate (*r = λ - µ*) and the turnover rate (ε) as it is easier to maximize the likelihood in this reparametrized likelihood landscape. For the BiSSE model, we used the MLE deployed under the R packages diversitree 0.9-3 (Fitzjohn 2012) and castor 1.6.6 (Louca and Doebeli 2018), both conditioned on the crown age of the phylogeny, under a Bernoulli sampling scheme and parametrized to infer *λ*_*1*_, *λ*_*2*_, *µ*_*1*_, *µ*_*2*_, and *q=q*_*12*_*=q*_*21*_. Diversitree is the traditional package used for fitting BiSSE; castor was developed more recently, and implements a faster algorithm for computing the likelihood of SSE models on large phylogenies (Louca and Doebeli 2018).

### PERFORMANCE ASSESSMENT

#### Accuracy of Parameter Estimation

To assess the accuracy of parameter estimation, we used 500 simulated test trees for the simple BD model and 10 000 for BiSSE. 17 BiSSE simulations for which castor and/or diversitree outputted an error message or no estimated values (15 for castor, 3 for diversitree with one simulation in common) were excluded from these analyses, resulting in 9.983 test trees.

To avoid over-penalizing the MLE approaches that, contrary to the NNs, do not have constrained parameter ranges, we added similar constraints to the MLE estimates. Indeed, to the exception of λ for constant-rate BD and λ_1_ for BiSSE for which NNs can predict values outside of the parameter values initially covered by the simulations due to tree rescaling, the other parameter estimates (ε, λ_2_ and q=q_12_=q_21_) are constrained by the parameter range used for ε, r_λ2_ and r_q_ in our simulations. We imposed similar constraints to the MLE estimates to prevent overpenalizing MLE in accuracy comparison. For example, if the MLE for ε is above 1, we set it to 1, and if the MLE for λ_2_ is lower than 0.1_*_λ_1_, we set it to 0.1*λ_1_ (0.1 is the minimum value for r_λ2_ used in our simulations).

To evaluate the distribution of the bias among the test set, we calculated the bias between the true (simulated, or ‘target’) parameter values and the predicted values per predictions as follows:

- *Bias* B_i_= predicted_i_ − target_i_

We computed three measures of error between the true parameter values and the predicted values on the test set:

- Mean absolute error 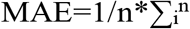 abs(predicted_i_ − target_i_)
- Mean relative absolute error 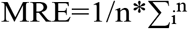 abs(predicted − target)/target
- Mean bias 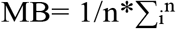 (predicted − target)

We also report the Pearson correlation coefficient between simulated and predicted values, computed with the R package ‘stats’ (‘cor.test’ function) version ‘4.2.1’.

Finally, we assessed the influence of tree size on parameter estimation accuracy.

#### Time Efficiency

We compared the average time of estimation between CNN-CDV and MLE for BiSSE. For the CNN-CDV approach, we reported the average CPU time of encoding a tree (averaged over 1,000,000 trees). The estimation on itself is negligible with respect to the time of encoding. For MLE BiSSE estimation with the castor and diversitree packages, we reported the average CPU time (average over 10.000 test trees, out of which 18 resulted in an error).

### EMPIRICAL ILLUSTRATION

Primates can have an antagonistic interaction with plants through herbivory, or a mutualistic one through frugivory and seed dispersal (Gómez and Verdú 2012). To illustrate the application of our deep learning approach, we reanalyzed the dataset of Gómez and Verdú (2012), that categorizes primate species according to the nature of their interaction with plants (state 1 for a mutualistic interaction, and 2 for an antagonistic one), using the primates phylogeny of Fabre et al. (2009). We started by pruning taxa without information on interaction type (13/273) and rescaled the phylogeny as previously described. Next, we performed some sanity checks to verify that the empirical data fell within the space covered by our BiSSE simulations; if it does not, this means that the model and/or the range of parameters used in the simulations is not well adapted to the empirical data, in which case application of the trained neural network to the data might output meaningless results. For these analyses, we used the set of phylogenies that we simulated under the BiSSE model to produce our test set. First, we checked that each of the summary statistics values calculated on the empirical data fell within the range spanned by the summary statistics values calculated on the simulated phylogenies. Then, we performed a principal component analysis (R package ‘FactoMineR’, function ‘PCA’) on the set of summary statistics described above (see Methods) with the addition of four simple summary statistics adapted to the inclusion of tips data: the number of tips in each state (1 or 2) and the phylogenetic diversity of each state (R package ‘picante’, function ‘pd’). Finally, we transformed the data into its CDV representation and fed it to the trained CNN-CDV network on the BiSSE model. We used a sampling fraction of 0.68, computed using a global diversity of 381 stable species complexes for primates, following Gómez and Verdú (2012).

## RESULTS

### PERFORMANCE ASSESSMENT

The comparison of parameter estimates obtained using deep NNs versus MLE for the BD model shows that CNN-CDV and FFNN-SS are as accurate as MLE, while FFNN-CDV has a lower accuracy, in terms of bias, absolute and relative error, for the speciation, extinction, net diversification and turnover rates (**Fig. 2, Table S1**). The likelihood of BD model has an exact analytical solution. This implies that MLE, together with CNN-CDV and FFNN-SS, are as accurate as one can be. The good performance of FFNN-SS might be explained by the representation of the lineage-through-time plot in the summary statistics, that contains all information available in the tree for homogeneous BD models (Nee et al. 1994). The lower performance of FFNN-CDV was expected given that FFNN is less adapted to unstructured data. The neural networks seem to avoid cases of high negative bias on the turnover rate compared to MLE: while this bias can reach down to −0.6 with MLE, it never falls behind −0.35 with CNN- CDV and FFNN-SS (**Fig. 2d**). This is probably due to the fact that contrary to MLE, CNN-CDV and FFNN-SS rarely return estimates close to 0 for the turnover rate (**Fig. S1d**).

**Figure 2:**
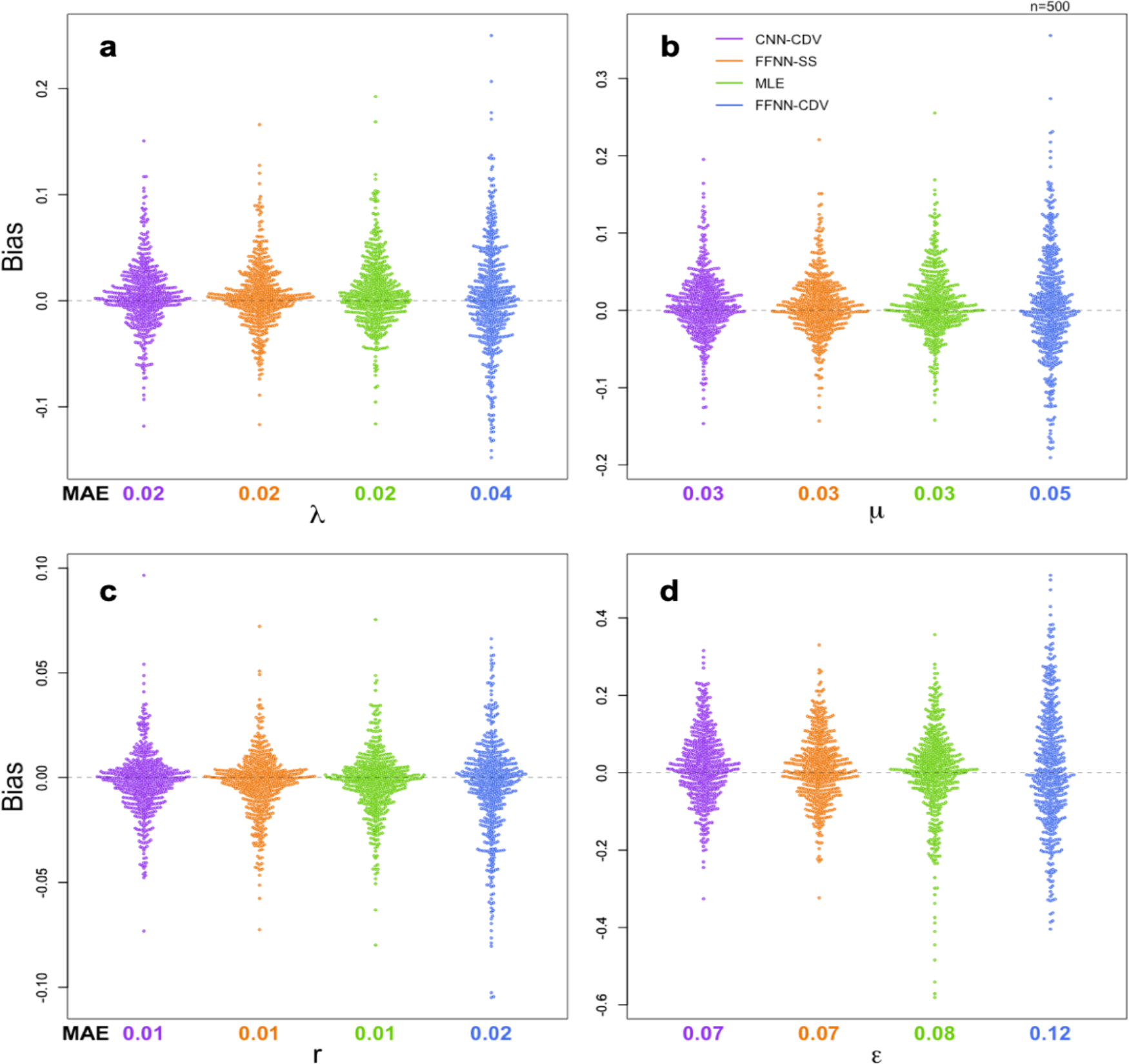
Comparison of estimation accuracy between pretrained CNN-CDV, FFNN-SS, FFNN-CDV and MLE for the BD model. Swarm plots representing the distribution of estimation biases across 500 simulations. Each dot represents the bias of a single simulation. Estimated parameters were obtained with Convolutional Neural Networks- Complete Diversity-reordered Vector (CNN-CDV in purple), Feed-Forward Neural Networks- Summary Statistics (FFNN-SS in orange), Maximum Likelihood Estimation (MLE in green) and Feed-Forward Neural Networks- Complete Diversity-reordered Vector (FFNN-CDV in blue) for (a) the speciation, (b) the extinction, (c) the net diversification and (d) the turnover rate. The mean absolute error (MAE) is displayed under each swarm plot.

The comparison of parameter estimates obtained using deep learning versus MLE for the BiSSE model confirms that CNN-CDV is at least as accurate as MLE. The only exception is for λ_2_ estimates, where MLE implemented in diversitree is slightly more accurate than CNN-CDV in terms of mean absolute error. CNN-CDV is more accurate than the fast MLE algorithm implemented in castor for all parameters (**Fig. 3, Table S2**). The accuracy of the MLE estimation implemented in castor is similar to that of the least reliable FFNN-CDV neural network; it performs slightly better for some parameters (*µ*_*1*_ and *q*_*12*_) but slightly worse for others (*µ*_*2*_). The CNN trained on a CDV without the individual tip information (CNN-CDV-less) is much less accurate, indicating that the information on individual tips states is well presented in the CDV representation, and extracted by the CNN (**Table S2**).

**Figure 3:**
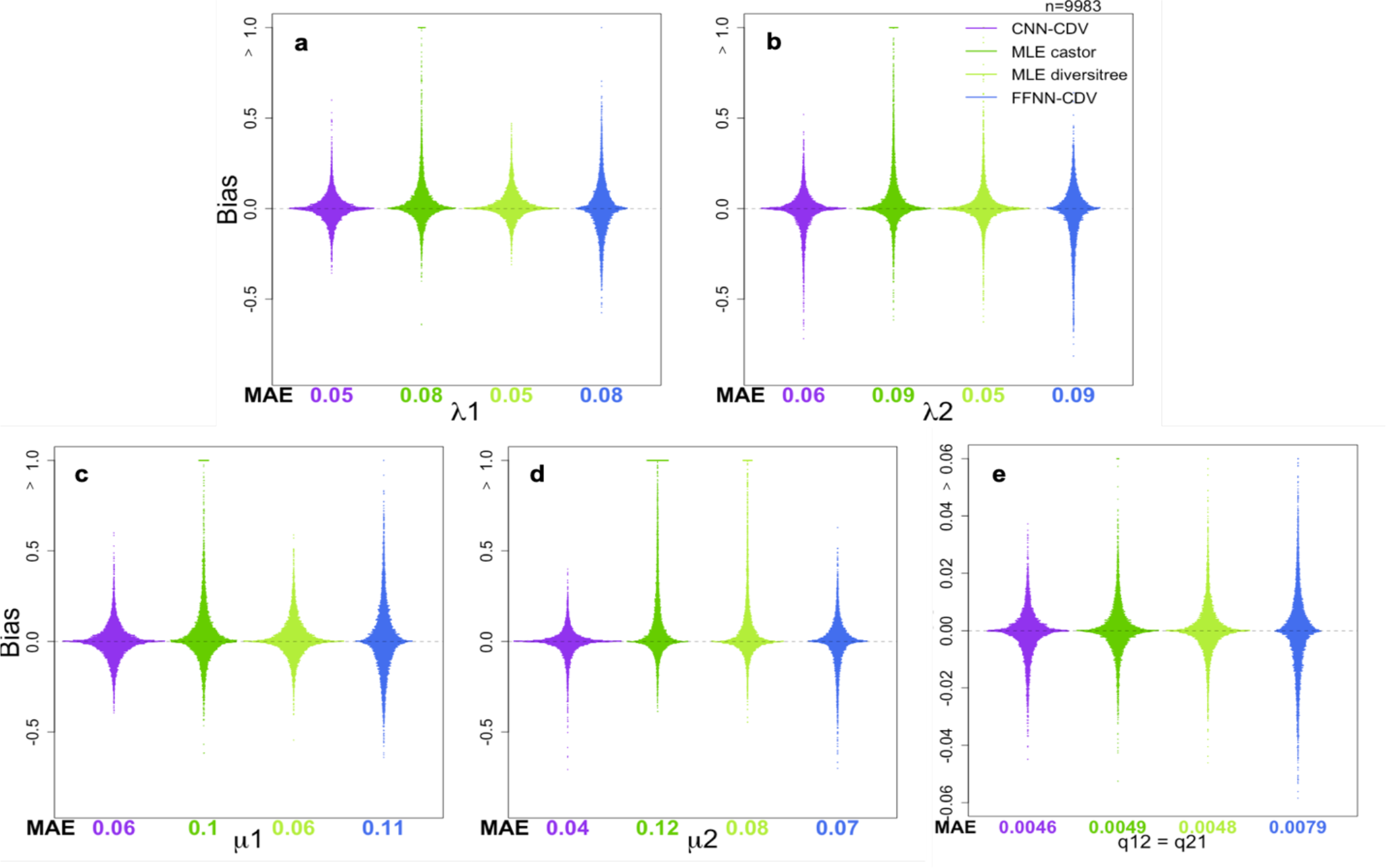
Comparison of estimation accuracy between pretrained CNN-CDV, FFNN-CDV, and MLE obtained with two different inference software for the BiSSE model. Swarm plots representing the distribution of estimation biases across 9,983 simulations. Estimated parameters were obtained with CNN-CDV (in purple), castor (in dark green), diversitree (in light green) and FFNN-CDV (in blue) for a) the speciation rates 1 and b) 2, c) the extinction rates 1 and d) 2 and e) the transition rates (q_*12*_*=q*_*21*_, in our setting). The simulations for which castor or diversitree outputted an error message (15/10,000 for castor and 3/10.000 for diversitree) were excluded from the comparison. The MAE is displayed under each swarm plot. For visualization purposes, the upper outliers of the swarm plot are not displayed at their extreme values but instead are put at the boundaries of the chart.

For both the BD and BiSSE model, the accuracy of CNN-CDV increases as the tree size increases (**Fig. 4**). Similar to MLE, a tree size of at least 300 sampled extant species is required for a median relative absolute error of 6% for the speciation rate and 18% for the extinction rate in the case of the BD model (**Fig. S3**). In the case of the BiSSE model, a tree size of at least 380 is required for a median relative absolute error of less than 14% on the speciation rates, 25% on the extinction rates, and 13% on the transition rate.

**Figure 4:**
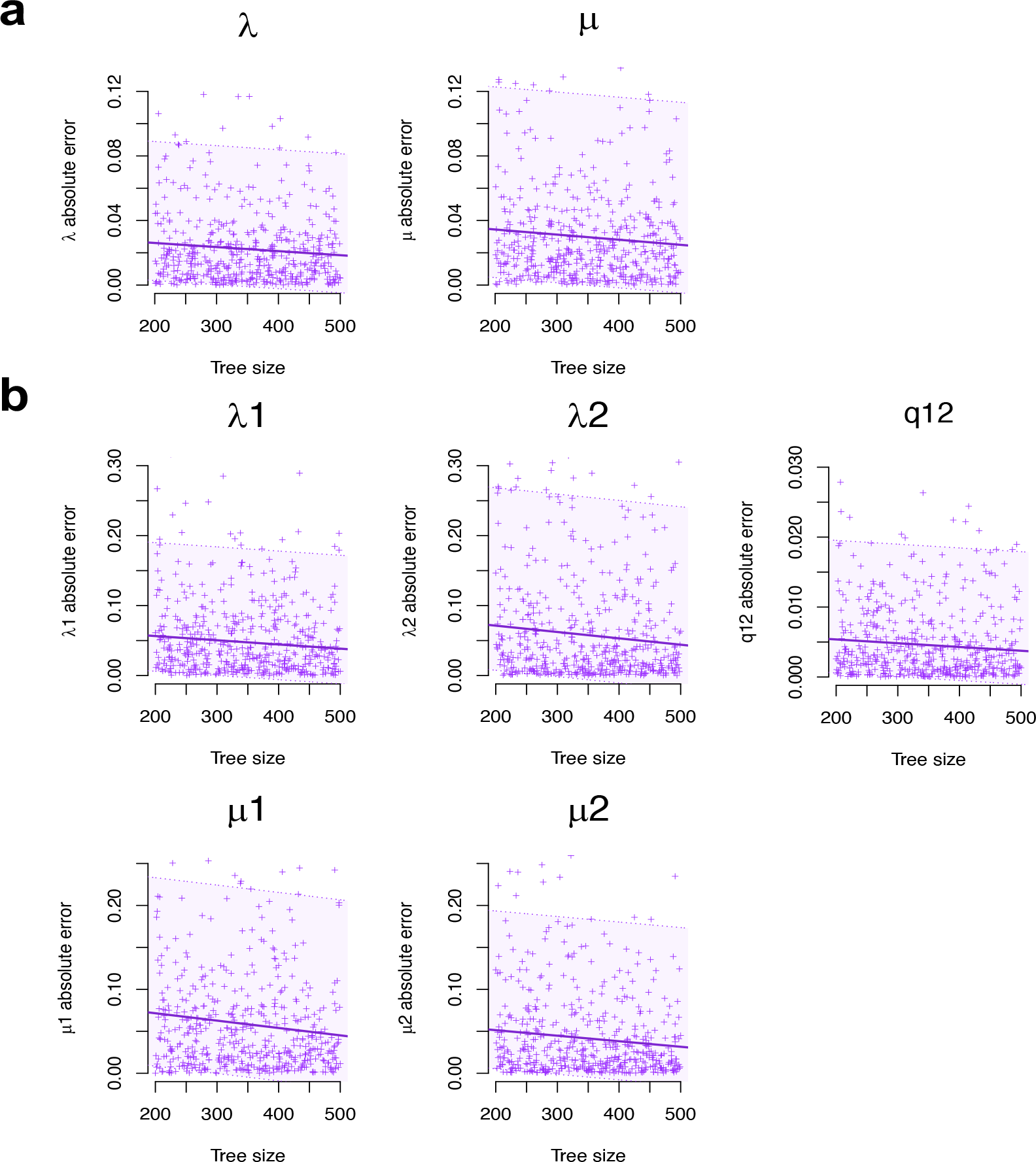
Effect of tree size on the absolute error of parameter estimates when using CNN- CDV under the BD and BiSSE models. For each model a) constant-rate birth-death (BD), and b) Binary-State Speciation and Extinction (BiSSE), we display the regression on absolute error for each parameter as a function of tree size for 500 test trees (for BiSSE 500 values are shown instead of 10.000 for visualization purposes). The area around the solid line delimited by the dotted lines represent the 95% confidence interval around the regression.

The parameter estimation for the 10.000 BiSSE simulations took 629.5 CPU hours with diversitree, 55.6 CPU hours with castor and 0.4 CPU hours with CNN-CDV, which consisted in encoding the test set into CDV. Once the CNN-CDV is trained, the estimation is thus around 140 times faster than castor and over 1500 times faster than diversitree. Training the network entailed first simulating the training set (1 million trees, 40 CPU hours). The training in itself then took 8 CPU hours. Contrary to MLE for which a large number of CPU hours are required for each new empirical analysis, with deep learning the empirical analyses are very fast once the network has been trained. The same pre-trained networks should be applicable to a large variety of empirical trees, thanks to tree rescaling which enables applications to clades of very different ages and speciation rates.

### EMPIRICAL ILLUSTRATION

The primate phylogeny with associated character state (mutualistic or antagonistic interaction with plants) passed the sanity checks on model adequacy. Indeed, all the summary statistics computed on the empirical data fell within the range spanned by the simulations. Likewise, our PCA analysis on these summary statistics showed the empirical data nested in the simulations space, when considering both PC1 and PC2 (explaining together 69% of the variance of our data, **Fig. 5a**) and PC3 and PC4 (that represent an additional 12 points of explained variance, **Fig. 5b**). The CNN-CDV analyses estimated a speciation rate of 0.295 for primate species with a mutualistic interaction with plants, and of 0.093 for those with an antagonistic interaction. The turnover rate ε was estimated at 0.234, and the transition rate at 0.0089. The resulting estimated net diversification rate is 0.225 for primates with a mutualistic interaction, and 0.071 for those with an antagonistic interaction. This simple analysis suggests that mutualistic interactions can favor diversification in primates, as found by (Gómez and Verdú 2012), although we do not interpret this result further here, as models with hidden traits should be used to reach more convincing biological conclusions (Beaulieu and O’Meara 2016). The goal of this empirical analysis is simply to illustrate how the method and trained networks can be used on empirical data.

**Figure 5:**
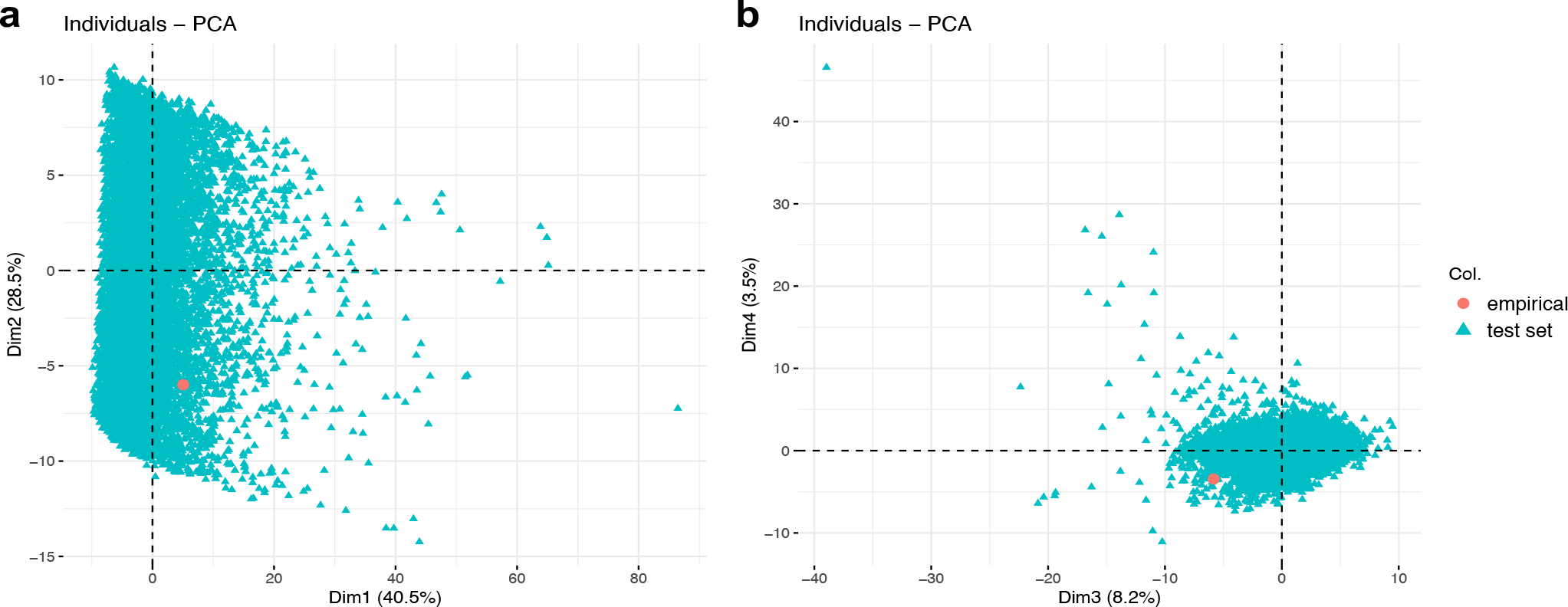
The primate data falls within the space of our BiSSE simulations. Coordinates of the empirical data (in pink) and of the test set simulations (turquoise) on the a) PC1 and PC2 axes and b) PC3 and PC4 axes of a principal component analysis performed on 102 summary statistics.

## DISCUSSION

We developed, tested and illustrated the use of a deep learning based inference approach for phylogenetic diversification analyses, including the case of trait-dependent diversification. We found that both convolutional neural networks combined with a compact representation of the phylogenetic data into a matrix (the CDV) and feed-forward neural networks combined with summary statistics can reach levels of parameter estimation accuracy comparable to those obtained with the well-established maximum likelihood approach, while being faster by several orders of magnitude.

To demonstrate the potential of deep learning for phylogenetic diversification analyses, we worked with two simple diversification models on which MLE estimates can easily be obtained for comparison (the BD and BiSSE models). We also worked with phylogenies of relatively moderate size (200 to 500 extant species sampled). The real value of deep learning will be to allow rapid inferences for more complex models for which likelihoods are not tractable or long to compute and for large phylogenies of several thousands of extant species. Our analyses on simple models demonstrate that it is worth putting efforts and computation power into simulating more complex models, generating larger trees, and training neural networks on such simulations. Given our results, we can expect efficient simulators combined with the CDV representation and CNN to provide an accurate likelihood-free estimation method applicable to very large phylogenies. The CDV representation is not model-specific and can easily be enriched with information on both internal nodes and tips. It could be used for example to represent information on species multidimensional traits, geographic distributions, abundances, and genetic diversity. Combined with efficient simulation models for the evolution of biodiversity (Hagen et al. 2021), the CNN-CDV deep learning inference approach could help adjusting biologically realistic biodiversity models to multifaceted data for a better understanding of how present-day biodiversity was generated, maintained, and distributed geographically.

Parameter estimates are more meaningful when associated with a measure of confidence. With our deep learning framework, obtaining a confidence interval around the estimates can be achieved through approximated parametric bootstrapping (Voznica et al. 2022), which uses the distribution of prediction error measured on simulations. Besides parameter estimation, diversification models are widely used for model comparison, in order to test alternative hypotheses about how diversification proceeds. This problem can be treated with deep learning as a classification problem, with neural networks trained to distinguish simulations from different models. This approach has been developed in Voznica et al. (2022) and shown to perform well. In the case of the BiSSE model for example, a neural network could be trained to distinguish data simulated under a model where states influence speciation rates from data simulated under a model where they do not. Given that speciation rates can be estimated with good accuracy with deep learning, we expect that neural networks will also be able to efficiently distinguish these models given enough differences in speciation rates between states.

We have explored the applicability of CNNs and FFNNs for phylogenetic diversification inference. Other classes of neural networks may also bring accurate solutions for parameter estimation and model selection when combined with simulated datasets. Noticeably, the Graph Neural Networks or Graph Convolutional Networks are neural networks designed for graph data that could be particularly well suited for analyzing phylogenies, encoded as directed graphs. Applications of the GNNs have been developed in the last couple of years across different fields, for instance in protein interaction prediction, drug design or social networks analyses (Zhou et al. 2020). More work is required to assess which network architecture performs best for phylogenetic diversification analyses, which has been initiated in Laajaiti et al..

As the ground truth is unknown, the neural networks are trained on simulations. This raises questions on how robust such approaches are to model misspecification, even though some studies suggest that machine learning approaches might be more robust to model misspecification than other inference approaches (Liang and Jordan 2008; Lee et al. 2010). We illustrated how some sanity checks can be performed to verify that the empirical data falls within the space of simulated data. If it does not, outputs of the neural networks should not be trusted. As in Voznica et al. (2022), we used summary statistics for these sanity checks. Another approach circumventing the use of summary statistics would consist in using autoencoders (Hinton and Salakhutdinov 2006), often deployed for anomaly detection (Chalapathy and Chawla 2019). The autoencoders are a family of NNs, which are trained to output their input while enforcing dimensionality reduction within their neural layers. The induced reconstruction error, *i*.*e*. the difference between the input and output, can then be used to check if an empirical data is well represented by simulations. For example, autoencoders could be trained on the CDV representation of phylogenetic data simulated under diversification models, and then applied to empirical phylogenetic data. If the reconstruction error is larger for the empirical data than for the simulations, this indicates departures from the simulated model.

We have considered the task of fitting birth-death diversification models to a fixed phylogeny, assumed to be known. While this is a current practice in the field, a better way to account for phylogenetic uncertainty consists in performing full phylogenetic inference, where the phylogenetic tree is inferred from molecular sequence alignments jointly with the parameters of the diversification process (Bouckaert et al. 2014, 2019). Machine learning is already used to data mine molecular sequence alignments (Yang et al. 2020), and recent progress has been made for phylogenetic reconstruction as well (Suvorov et al. 2020; Zou et al. 2020; Nesterenko et al. 2022; Solis-Lemus et al. 2022). Ultimately, these recent advances could be in the long term combined to train CNNs directly on sequences simulated from a joint process of speciation, extinction, and sequence evolution.

Deep learning is gaining popularity in biology, including in ecology and evolution (Borowiec et al. 2021). It has been used as a likelihood-free approach to fit population genetic models (Flagel et al. 2019) to sequence data, and more recently to fit epidemiological models to pathogen phylogenies (Voznica et al. 2022). We have shown that it can also perform well as a likelihood-free approach for fitting diversification models to phylogenies of extant species. More work is needed to establish which data representation and network architecture perform best, to perform statistical inference directly on sequence alignments rather than on fixed phylogenies, and to efficiently train networks for more complex models. We hope that our paper will stimulate research in this direction. Ultimately, this should foster the development of new diversification models that are not limited (or whose design is not biased) by our ability to compute likelihoods.

## ACKNOWLEDGEMENTS

SL was supported by PSL IRIS Science des données, données de la science and the Fondation pour la Recherche Médicale (FDT202106013269). JV was supported by the Ecole Normale Supérieure Paris-Saclay and by ED Frontières de l’Innovation en Recherche et Education, Programme Bettencourt. HM acknowledges funding from ERC-CoG grant PANDA. We thank Quang Tru Huynh for administrating the GPU farm at Institut Pasteur and the INCEPTION program (Investissement d’Avenir grant ANR16-CONV-0005) that financed the GPU farm.

## DATA AVAILABILITY

All data and codes underlying this article are available on GitHub, at https://github.com/JakubVoz/deeptimelearning [doi].

