## Supplementary Figures and Tables for "Deep Learning from Phylogenies for Diversification Analyses"

**SUPPLEMENTARY INFORMATION**

**SUPPLEMENTARY TABLE 1: Comparison of accuracy and precision for machine learning and maximum likelihood estimation techniques for constant-rate Birth Death model.**

| Parameters | Mean Absolute Error | | | | Mean Relative Error | | | |
|---|---|---|---|---|---|---|---|---|
| | MLE | FFNN-Sumstats | CNN-CDV | FFNN-CDV | MLE | FFNN-Sumstats | CNN-CDV | FFNN-CDV |
| **Speciation rate** | 0.02 | 0.02 | 0.02 | 0.04 | 0.09 | 0.08 | 0.08 | 0.14 |
| **Extinction rate** | 0.03 | 0.03 | 0.03 | 0.05 | 0.51 | 0.62 | 0.62 | 1.18 |
| **Net diversification rate** | 0.01 | 0.01 | 0.01 | 0.02 | 0.11 | 0.10 | 0.11 | 0.17 |
| **Turnover rate** | 0.08 | 0.07 | 0.07 | 0.12 | 0.40 | 0.48 | 0.50 | 0.88 |
| **Parameters** | **Pearson Correlation Coefficient** | | | | **Mean Bias** | | | |
| | MLE | FFNN-Sumstats | CNN-CDV | FFNN-CDV | MLE | FFNN-Sumstats | CNN-CDV | FFNN-CDV |
| **Speciation rate** | 0.97 | 0.98 | 0.97 | 0.93 | 0.008 | 0.005 | 0.005 | 0.003 |
| **Extinction rate** | 0.93 | 0.93 | 0.93 | 0.78 | 0.010 | 0.007 | 0.008 | 0.006 |
| **Net diversification rate** | 0.99 | 0.99 | 0.99 | 0.98 | -0.002 | -0.003 | -0.002 | -0.005 |
| **Turnover rate** | 0.92 | 0.95 | 0.94 | 0.82 | 0.0008 | 0.013 | 0.019 | 0.015 |

*For each estimated parameter and estimation method, we evaluated the accuracy on 500 test simulations in terms of: 1] Mean Absolute Error (MAE) in green, 2] Mean Relative Error (MRE) in blue, 3] Pearson correlation coefficient in yellow and 4] Mean Bias in grey. The compared estimation methods are a) Maximum likelihood estimation MLE-based method, b) Feed-Forward Neural Network trained on summary statistics (FFNN-SS), c) Convolutional Neural Network trained on the CDV representation (CNN-CDV), d) FFNN trained on the CDV representation (FFNN-CDV). The MRE for turnover was very high with each method, which is partially due to the covered parameter space (uniformly covered between 0.01 and 1). The CNN-CDV exhibits a higher bias for the turnover rate.*

**SUPPLEMENTARY TABLE 2: Comparison of accuracy and precision for machine learning and maximum likelihood estimation techniques for Binary State Speciation and Extinction model.**

| Parameters | Mean Absolute Error | | | | | Mean Relative Error | | | | |
|---|---|---|---|---|---|---|---|---|---|---|
| | castor | diversitree | CNN-CDV | CNN-CDV-less | FFNN-CDV | castor | diversitree | CNN-CDV | CNN-CDV-less | FFNN-CDV |
| **Speciation rate 1** | 0.076 | 0.048 | 0.047 | 0.150 | 0.084 | 0.16 | 0.10 | 0.10 | 0.31 | 0.17 |
| **Speciation rate 2** | 0.086 | 0.055 | 0.058 | 0.110 | 0.085 | 0.42 | 0.22 | 0.24 | 0.42 | 0.38 |
| **Extinction rate 1** | 0.097 | 0.064 | 0.058 | 0.146 | 0.109 | 3.15 | 1.65 | 1.80 | 4.58 | 4.08 |
| **Extinction rate 2** | 0.116 | 0.079 | 0.041 | 0.089 | 0.067 | 12.38 | 3.06 | 1.94 | 4.44 | 4.27 |
| **Transition rate** | 0.005 | 0.005 | 0.005 | 0.010 | 0.008 | 0.21 | 0.21 | 0.19 | 0.46 | 0.36 |

| Parameters | Pearson Correlation Coefficient | | | | | Mean Bias | | | | |
|---|---|---|---|---|---|---|---|---|---|---|
| | MLE | diversitree | CNN-CDV | CNN-CDV-less | FFNN-CDV | castor | diversitree | CNN-CDV | CNN-CDV-less | FFNN-CDV |
| **Speciation rate 1** | 0.77 | 0.97 | 0.97 | 0.74 | 0.92 | 0.038 | 0.017 | 5e-04 | 0.003 | 6e-04 |
| **Speciation rate 2** | 0.67 | 0.93 | 0.92 | 0.75 | 0.84 | 0.045 | 0.012 | -0.010 | -0.013 | -0.016 |
| **Extinction rate 1** | 0.65 | 0.92 | 0.92 | 0.48 | 0.73 | 0.040 | 0.016 | -0.002 | 1e-04 | 0.009 |
| **Extinction rate 2** | 0.44 | 0.74 | 0.91 | 0.57 | 0.78 | 0.083 | 0.042 | -0.006 | -0.003 | -0.003 |
| **Transition rate** | 0.94 | 0.94 | 0.95 | 0.71 | 0.85 | 5e-04 | 5e-04 | -8e-04 | 8e-04 | -4e-04 |

*For each estimated parameter and estimation method, we evaluated the accuracy on 9.983 test simulations in terms of: 1] Mean Absolute Error (MAE) in green, 2] Mean Relative Error (MRE) in blue, 3] Pearson correlation coefficient in yellow and 4] Mean Bias in grey. The compared estimation methods are a) Maximum likelihood estimation MLE-based method using the R package castor, b) MLE-based method using the R package diversitree, c) Convolutional Neural Network trained on the CDV representation (CNN-CDV), d) CNN trained on CDV-less without individual information on tip state (CNN-CDV-less) and e) Feed-Forward Neural Network trained on the CDV representation (FFNN-CDV). The MRE for the turnover rate was very high for each method, which is partially due to the covered parameter space (uniformly covered between 0 and 1). With CNN-CDV, we obtain comparable results as with castor and diversitree.*

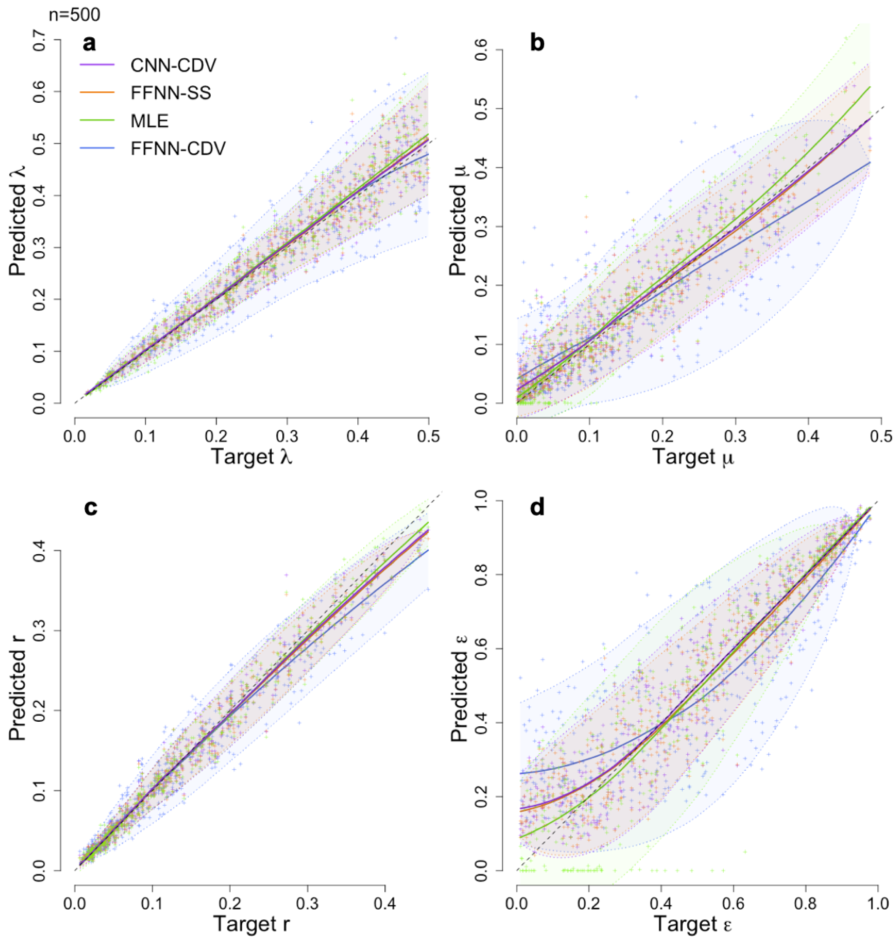


**SUPPLEMENTARY FIGURE 1**: **Predicted versus target plots.**

For each parameter predicted for BD we show a plot of predicted versus target values for 500 simulations from the test set, each dot representing one prediction: a) speciation rate λ, b) extinction rate $\mu$, c) net diversification rate r, d) turnover rate ε. The dashed line represents the identity line, while the full line is the local polynomial regression with its 95% confidence interval delimited by the dotted lines.

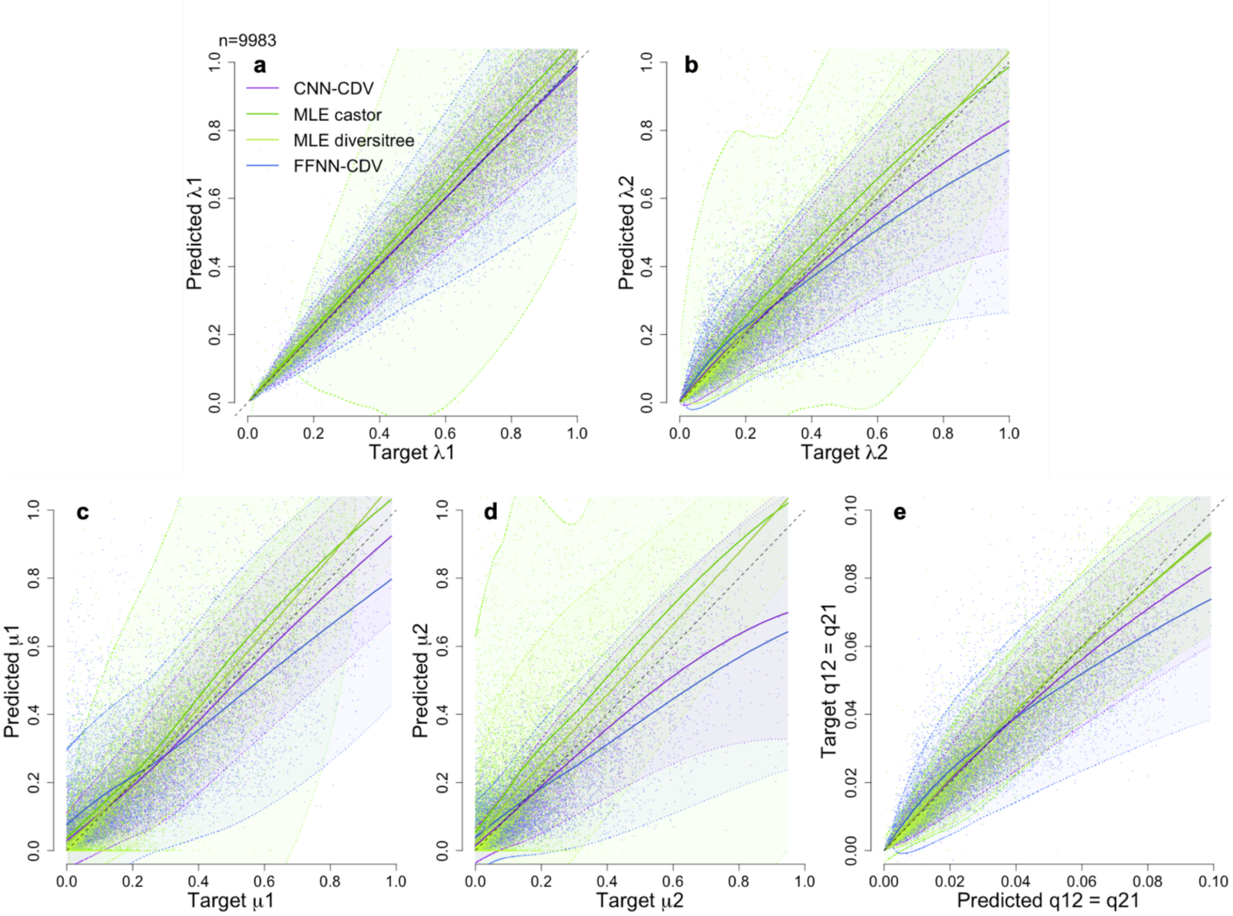


**SUPPLEMENTARY FIGURE 2** : **Predicted versus target plots.**

For each parameter predicted for BiSSE we show a plot of predicted versus target values for 9,983 simulations from the test set, each dot representing one prediction: a) speciation rate 1 $\lambda_1$, b) speciation rate 2 $\lambda_2$, c) extinction rate 1 $\mu_1$, d) extinction rate 2 $\mu_2$, e) the transition rates $q_{12}=q_{21}$. The dashed line represents the identity line, while the full line is the local polynomial regression with its 95% confidence interval delimited by the dotted lines.

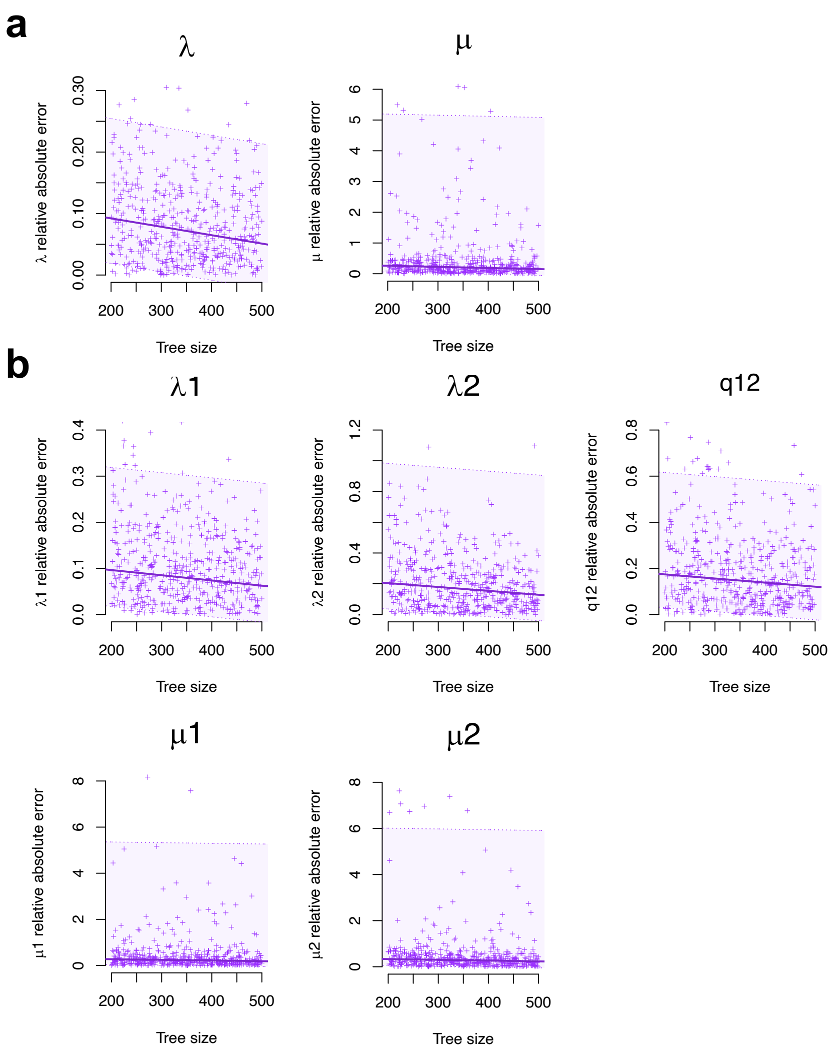


**SUPPLEMENTARY FIGURE 3** : ***Relative accuracy of CNN-CDV increases with the tree size.***

*For each model a) constant-rate BD, and b) BiSSE, we display the quantile regression on the relative absolute error in solid line for each parameter as a function of tree size for 500 test trees (for BiSSE, 500 values are shown instead of 10.000 for visualization purposes). The area around the solid line delimited by the dotted lines represent the 95% confidence interval around the quantile regression.*
